# RIP3 and MLKL regulate Hepatic ER stress in alcohol-associated liver disease and pharmacological ER stress models: insights beyond necroptosis

**DOI:** 10.1101/2025.09.24.677754

**Authors:** Rakesh K Arya, Emily Huang, Megan R McMullen, Kyle L Poulsen, Jianguo Wu, Jared Travers, Evi Paouri, Dimitrios Davalos, Laura E Nagy

**Affiliations:** Northern Ohio Alcohol Center, Department of Inflammation and Immunity, Cleveland Clinic, Cleveland OH, United States; Department of Pharmacology and Toxicology, College of Osteopathic Medicine, Michigan State University, East Lansing, MI, United States; Department of Molecular Medicine, Case Western Reserve University, Cleveland, OH, United States; Department of Gastroenterology and Liver Disease, University Hospitals Cleveland Medical Center, Cleveland, OH, United States; Department of Neurosciences, Lerner Research Institute, Cleveland Clinic, Cleveland, OH, United States; Department of Gastroenterology and Hepatology, Cleveland Clinic, Cleveland, OH, United States

**Keywords:** RIP3, RIP3 kinase, MLKL, ER stress, ER expansion, Cell death, alcohol-associated liver disease

## Abstract

**Background and Aim:** Endoplasmic reticulum (ER) stress is an important contributor to liver disease progression, including alcohol-associated liver disease (ALD). While receptor-interacting protein kinase-3 (RIP3) and mixed lineage kinase domain-like pseudokinase (MLKL) are known for their roles in necroptosis, emerging evidence highlights their non-canonical functions in metabolic regulation and cellular stress responses. However, their specific role in regulating hepatic ER stress remains unclear. This study investigates how RIP3, its kinase activity, and MLKL regulate ER stress pathways during chronic ethanol exposure and pharmacological ER stress induction.

**Methods:** *Rip3^-/-^*, *Rip3^K51A/K51A^* and *Mlkl^-/-^* mice alongside WT controls and pharmacological necroptosis inhibitors were used to study the role of RIP3 and MLKL in modulating ER stress. Chronic ethanol feeding and pharmacological agents (tunicamycin, thapsigargin) were utilized to induce ER stress *in vivo* and in isolated primary hepatocytes. ER stress markers were assessed by qPCR and western blot, ER expansion was evaluated by confocal microscopy, and hepatocyte viability was measured using MTS assay.

**Results:** Chronic ethanol increased expression of ER stress markers in WT mice; this response was attenuated in *Rip3^-/-^* mice. Tunicamycin exposure increased hepatic ER stress markers in WT mice; this response was diminished in *Rip3^-/-^*, *Rip3^K51A/K51A^* and *Mlkl^-/-^*mice. In primary hepatocytes, genetic and pharmacological inhibition of RIP3 and MLKL also reduced thapsigargin-induced ER stress responses. Hepatocytes isolated from *Rip3^-/-^*, *Rip3^K51A/K51A^*and *Mlkl^-/-^* mice exhibited enhanced cell viability under ER stress conditions compared to hepatocytes from WT mice, which was associated with ER expansion as a potential mechanism for mitigating ER stress.

**Conclusion:** This study highlights a novel function of RIP3 and MLKL in regulating hepatic ER stress responses, expanding their known roles beyond programmed necrosis.

**Impact and Implications:** This study provides new mechanistic insight into how RIP3 and MLKL regulate hepatic ER stress responses, extending their roles beyond necroptosis. By demonstrating that genetic or pharmacological inhibition of *Rip3*, RIP3 kinase activity and *Mlkl* attenuates ER stress signaling, reduces cell death, and promotes adaptive ER remodeling, our findings identify these proteins as key modulators of hepatocyte survival under stress. These results are important for researchers and clinicians focused on alcohol-associated liver disease and other ER stress–driven liver disorders, as they highlight novel therapeutic targets. In practical terms, modulation of the RIP3– MLKL axis could inform the development of interventions aimed at enhancing ER stress resilience, with potential applications in drug development for ER stress–associated liver injury.

## Introduction

The liver plays a central role in maintaining metabolic homeostasis through regulation of glucose and lipid metabolism, detoxification, and protein synthesis, making it critical for overall metabolic health ^[1]^. Hepatocytes, the predominant cell type in the liver, contain an extensive endoplasmic reticulum (ER) network essential for these functions ^[2, 3]^. However, chronic alcohol consumption or metabolic stress can disrupt normal ER function, leading to ER stress-associated toxicity and contributing to various liver disorders. Among these, alcohol-associated liver disease (ALD) has emerged as a significant health concern, encompassing a spectrum of conditions from hepatic steatosis to inflammation and progression to fibrosis or cirrhosis ^[4–6]^. Other related conditions include metabolic dysfunction-associated steatotic liver disease (MASLD) and viral hepatitis, all of which share ER stress as a common pathogenic mechanism ^[7, 8]^.

The ER is equipped with an adaptive response system called the unfolded protein response (UPR), initiated by three ER transmembrane protein sensors: Inositol-requiring enzyme 1 alpha (IRE1α), protein kinase RNA-like ER kinase (PERK), and activating transcription factor 6 (ATF6). Upon activation, IRE1α, with its dual endoribonuclease and kinase activities, generates the active transcription factor X-box binding protein 1 (sXBP1) through mRNA splicing, which upregulates genes involved in protein folding, secretion, and degradation of misfolded proteins. PERK activation leads to the selective translation of activating transcription factor 4 (ATF4), which in turn induces the expression of C/EBP homologous protein (CHOP), a key mediator of ER stress-induced apoptosis. ATF6, once activated, is cleaved and translocates to the nucleus to function as a transcription factor, stimulating the expression of ER chaperones such as GRP78 to enhance protein folding capacity. These three pathways work together to alleviate ER stress and restore cellular homeostasis. However, if ER stress is prolonged or unresolved, the UPR shifts from an adaptive to a maladaptive response, promoting cell death through mechanisms such as apoptosis or necroptosis ^[2, 9, 10]^.

Necroptosis is a regulated form of necrotic cell death mediated by two critical proteins: receptor-interacting protein kinase 3 (RIP3) and mixed lineage kinase domain-like pseudokinase (MLKL) ^[11]^. While the classical necroptotic pathway leads to MLKL-driven plasma membrane rupture and release of intracellular contents, growing evidence has revealed non-canonical functions of both RIP3 and MLKL beyond cell death. Recent studies indicate that RIP3 and MLKL can modulate diverse cellular processes, including the regulation of mitochondrial fission, inflammation, phagocytosis, and intracellular trafficking ^[12–15]^. Notably, MLKL has been found to localize not only to the plasma membrane but also to several intracellular organelles such as the mitochondria, ER, and lysosomes, where it may influence organelle function and contribute to various pathophysiological responses ^[16–18]^. These emerging roles suggest that RIP3 and MLKL act as broader regulators of cell stress and homeostasis, extending their influence well beyond their canonical roles in necroptotic cell death ^[19–21]^.

Despite these insights, the specific role of RIP3 and MLKL in regulating hepatic ER stress remains poorly understood, especially under stress conditions such as those induced by ethanol exposure or tunicamycin/thapsigargin, known ER stress inducers. Furthermore, while the kinase activity of RIP3 is typically required for MLKL activation and necroptosis, recent studies have pointed to kinase-independent functions of RIP3 in stress regulation and inflammation ^[15, 22, 23]^. This raises crucial questions about the relative contributions of RIP3 kinase domain in modulating ER stress responses. In this study, we aim to investigate the role of RIP3, its kinase domain, and MLKL in the regulation of hepatic ER stress and cell death. Using genetically modified mouse models (*Rip3^-/-^, Rip3^K51A/K51A^, and Mlkl^-/-^*) and pharmacological inhibitors, we assessed their contribution to ER stress signaling and adaptive mechanisms under ethanol- and tunicamycin/thapsigargin-induced stress conditions. Through exploring the crosstalk between RIP3-MLKL signaling and ER stress pathways, this study aims to enhance our understanding of the molecular mechanisms underlying ALD and identify potential therapeutic targets for ER stress-associated liver diseases.

## Materials and Methods

### Animal models

*Rip3^-/-^* mice were a generous gift from Dr. Vishva M. Dixit, (Molecular Oncology Department, Genentech, Inc., San Fransisco, CA). *Rip3^K51A/K51A^* mice were originally generated by GlaxoSmithKline (strain is viable and fertile). Specifically, this strain was created through a homologous recombination process, employing a targeting construct designed to introduce mutations that replaced the catalytic lysine residue with alanine (K51A). This modification effectively resulted in the complete abolition of RIP3 kinase activity ^[24]^. *Mlkl^−/−^* mice were purchased from Taconic Biosciences (TF2780). All mutant mice strains were either on C57BL/6J or backcrossed to a C57BL/6J background. WT controls were either C57BL/6J purchased from Jackson Laboratory (Bar Harbor, ME) or WT littermates. Animals were housed 2 per cage in standard micro isolator cages and maintained in a temperature-regulated facility with a 10h:14h light/dark cycle. All procedures using animals were approved by the Cleveland Clinic Institutional Animal Care and Use Committee.

### Chronic ethanol feeding

For the chronic ethanol-feeding models, 8- to 10-week-old female *Rip3^-/-^*mice and their wild-type littermate controls were randomized into ethanol fed and pair fed groups. The mice were initially acclimated to a control liquid diet for 2 days. The ethanol fed groups were given access to the diet containing ethanol with a progressive increase in ethanol concentration: 1% (vol/vol) for 2 days, 2% for 2 days, 4% for 1 week, 5% for 1 week, and finally 6% for the last week, totaling 25 days and reaching 32% of total calories. Control mice were pair-fed a control diet which iso calorically replaced maltose dextrins for ethanol. Body weights were recorded twice weekly and food consumption per cage was measured daily. At euthanasia, livers were perfused with saline, and portions of liver tissue were flash-frozen in liquid nitrogen and stored at -80°C for RNA or protein analysis or fixed in 10% formalin for histology. Blood was collected in EDTA-containing tubes to obtain plasma, which was then stored at -80°C.

### Tunicamycin treatment

8- to 10-week-old male and female WT mice or *Rip3^-/-^*or *Rip3^K51A/K51A^* or *Mlkl^-/-^* mice had *ad libitum* access to standard chow diet and water. Mice were randomly allocated into two groups based on their average body weight to achieve a balanced distribution of body weight within each group. Mice were challenged with a single intraperitoneal injection of vehicle (150 mM dextrose solution) or tunicamycin (0.5 mg/kg; T7765, Sigma Aldrich) dissolved in vehicle. Mice were euthanized at various time points post-injection as indicated in the figure legends, and liver and blood samples were collected. Tissue processing and storage were performed as described above. Additional materials and methods are provided in the Supplemental Information.

### Statistical analysis

All values in the figures are presented as means ± SEM. Statistical analysis was conducted using the general linear model procedure (SAS, Carey, IN). Data was log-transformed when necessary to achieve a normal distribution. Post hoc comparisons were performed using least square means testing. A p-value of <0.05 was considered statistically significant, and values with different superscripts indicate significant differences (*P* < 0.05).

## Results

### Activation of ER stress response pathways in patients with severe alcohol-associated hepatitis (sAH)

We performed GSEA on two independent bulk RNA-seq datasets from healthy individuals and patients with sAH undergoing liver transplant to determine if ER stress markers are dysregulated in sAH patient livers. Our analysis of 137 "endoplasmic"-related human gene sets from the GSEA-molecular signature database revealed significant enrichment of genes involved in the “positive_regulation_of_response_to_endoplasmic_reticulum_stress” in sAH patient livers (GSE143318 and DbGaP_phs001807) (**Fig. 1A and B**). These results indicate significant activation of ER stress response pathways in sAH patients, suggesting their potential role in ALD pathogenesis and as therapeutic targets.

**Figure 1.**
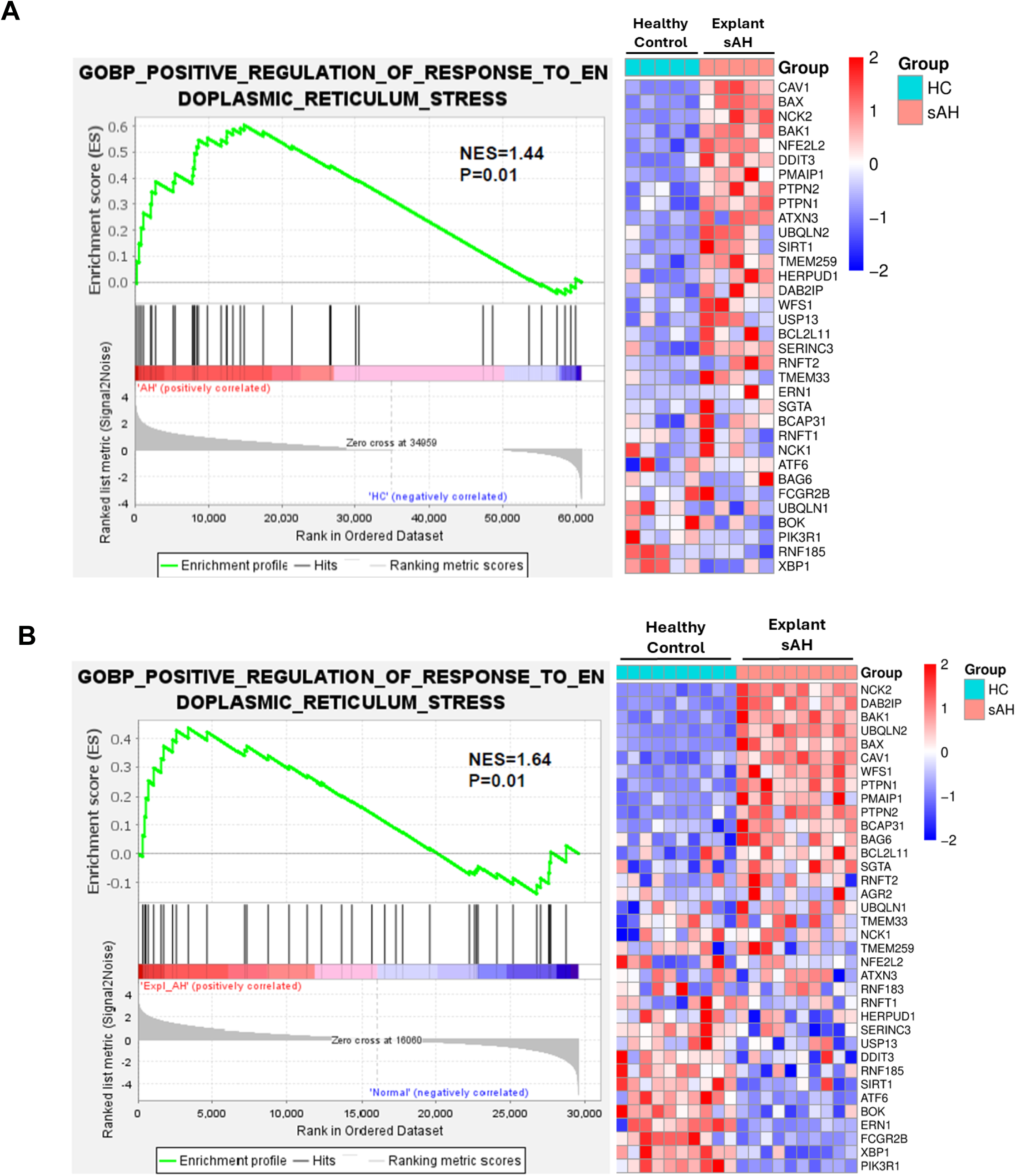
RNA-seq analyses reveal enrichment of genes positively responding to ER stress in severe alcohol-associated hepatitis liver samples. GSEA of two independent RNA-seq datasets comparing healthy controls (HC) and patients with sAH revealed upregulation of genes associated with the response to ER stress in sAH. A. GSEA analysis of dataset GSE143318: Bulk RNA-seq data from liver samples of healthy controls (n = 5) and explanted livers from patients with sAH (n = 5). B. GSEA analysis of DbGaP_phs001807: Bulk RNA-seq data from healthy controls (n = 10) and explanted livers from patients with sAH (n = 10). Heatmaps display log fold-change values for ER stress–associated genes, ranging from –2 (blue, downregulated) to +2 (red, upregulated).

### RIP3 contributes to ethanol-induced hepatic ER stress

Since ER stress is associated with the development of chronic ethanol-induced liver injury ^[2, 3, 7, 25]^ and *Rip3* deficient mice are protected from ethanol-induced liver injury ^[26, 27]^ (**Fig. S2A**). We tested the hypothesis that *Rip3* is required for the injurious hepatic ER stress response to ethanol. Chronic ethanol feeding to C57BL6/J WT mice increased expression of ER stress-related markers in the liver compared to littermate WT control mice, including increased expression of *sXbp1*, *Grp78* and C*hop* mRNA. In contrast, induction of *sXbp1, Grp78* and *Chop* mRNA were reduced in *Rip3^-/-^*mice in response to chronic ethanol. Expression of *Atf4* mRNA was not increased in either genotype after chronic ethanol feeding (**Fig. 2A**). At the protein level, ethanol feeding increased the expression of phospho-PERK and CHOP but not GRP78 in WT livers. In *Rip3*^-/-^ mice, the ethanol-induced upregulation of phospho-PERK and CHOP protein was prevented, although there was no reduction in GRP78 protein (**Fig. 2B**). Taken together, these data indicate that RIP3 contributes to chronic ethanol-induced ER stress and subsequent liver injury, potentially through PERK-CHOP-mediated pathway.

**Figure 2.**
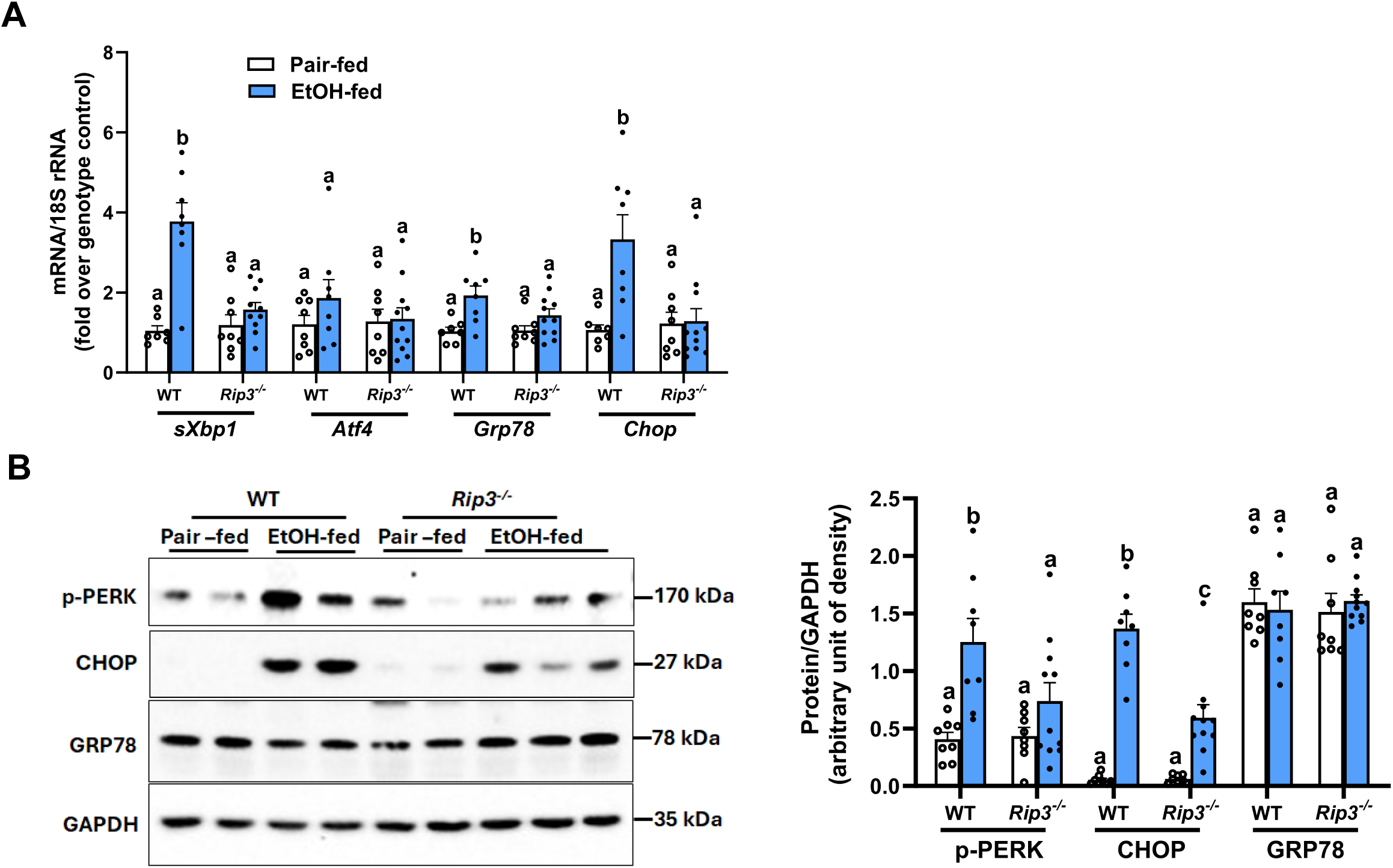
Deficiency of *Rip3* attenuates chronic ethanol-induced ER stress markers. WT and *Rip3^-/-^* mice were allowed free access to diets with increasing concentrations of ethanol (final concentration 32% of kcal) or pair-fed (control diet) for 25 days, as described in Methods. **A.** The mRNA expression levels of *sXbp1*, *Atf4*, *Grp78*, and *Chop* were measured by qRT-PCR and normalized to 18S rRNA. **B.** Relative expressions of phospho-PERK, GRP78, and CHOP were assessed by western blot, densities were quantified using ImageJ and normalized to GAPDH; n=8-11. Blots are processed in parallel for samples from the same experiment. Cropped blots are shown for clarity. Original blots are provided in Supplementary file “*Original western blot images*”. Values represent mean ± SEM. Values with different alphabetical superscripts are significantly different from each other. p < 0.05, assessed by ANOVA.

### ER stress-induced upregulation of RIP3 and MLKL in liver tissue

To elucidate the specific role of RIP3 in regulating ER stress, independent of ethanol’s multifaceted effects, we utilized tunicamycin, a well-established pharmacological inducer of ER stress. Tunicamycin inhibits N-linked glycosylation, crucial for protein folding, leading to misfolded protein accumulation in the ER and subsequent ER stress. Following tunicamycin challenge, we observed a time-dependent upregulation of ER stress-related proteins in the WT liver, including phospho-PERK, GRP78, and CHOP, with CHOP expression peaking at 18 hours (**Fig. S1A**). These responses were paralleled by time-dependent increases in plasma ALT (**Fig. S1B**). Notably, while RIP3 is typically expressed at low levels in healthy livers, it is induced by various stressors. Our previous work and that of others demonstrated that chronic ethanol upregulates expression of both RIP3 and ER stress markers in livers of mice and humans ^[26–29]^. In response to tunicamycin challenge, both RIP3 and MLKL proteins were increased within 6 hours of exposure to tunicamycin (**Fig. S1C**). To further define the cellular localization of these proteins, we performed immunohistochemistry (IHC) on liver sections. Based on morphologic assessment, tunicamycin markedly increased accumulation of RIP3 and phosphorylated MLKL (pMLKL), the activated form of MLKL, primarily in hepatocytes (**Fig. S1 D, E**). These findings indicate that a pharmacological inducer of ER stress increases expression of necroptotic mediators, independent of ethanol exposure, suggesting a potential mechanistic link between ER stress and RIP3-MLKL axis in liver injury.

### Tunicamycin challenge only induced modest hepatocellular injury

To investigate the contribution of RIP3 and MLKL in ER stress–mediated hepatocellular injury, liver histology, plasma ALT, and hepatic triglyceride concentrations were assessed in response to challenge with 0.5 mg/kg tunicamycin for 18 hours (**Fig. S2B-H**). H&E staining of liver sections revealed no overt morphological abnormalities across genotypes at this dose and time point (**Fig. S2 B, C and F**). Plasma ALT concentrations were modestly increased in response to tunicamycin in all genotypes. However, increases were lower in *Rip3^-/-^* and *Mlkl^-/-^* mice relative to WT controls, whereas ALT concentrations in *Rip3^K51A/K51A^*were between that of *Rip3^-/-^*and WT mice (**Fig. S2 D, G**). Hepatic triglyceride concentrations were increased in response to tunicamycin, independent of genotype (**Fig. S2 E, H**). These results indicate that the 18 hours time point tunicamycin induced only very mild liver injury and that genetic ablation of *Rip3* or *Mlkl* conferred partial protection, as reflected by reduced ALT release) and upregulated phospho-PERK, GRP78, and CHOP protein expression (**Fig. 3B-D**). However, *Rip3^-/-^* mice exhibited reduced expression of *sXbp1* and *Atf4* mRNA and lower levels of phospho-PERK, GRP78, and CHOP proteins. Notably, ATF6 protein expression was increased in response to tunicamycin independent of genotype (**Fig. 3A and 4B**). These results suggest that *Rip3* may primarily regulate the PERK and IRE1α (*sXbp1*) arms of the UPR.

**Figure 3.**
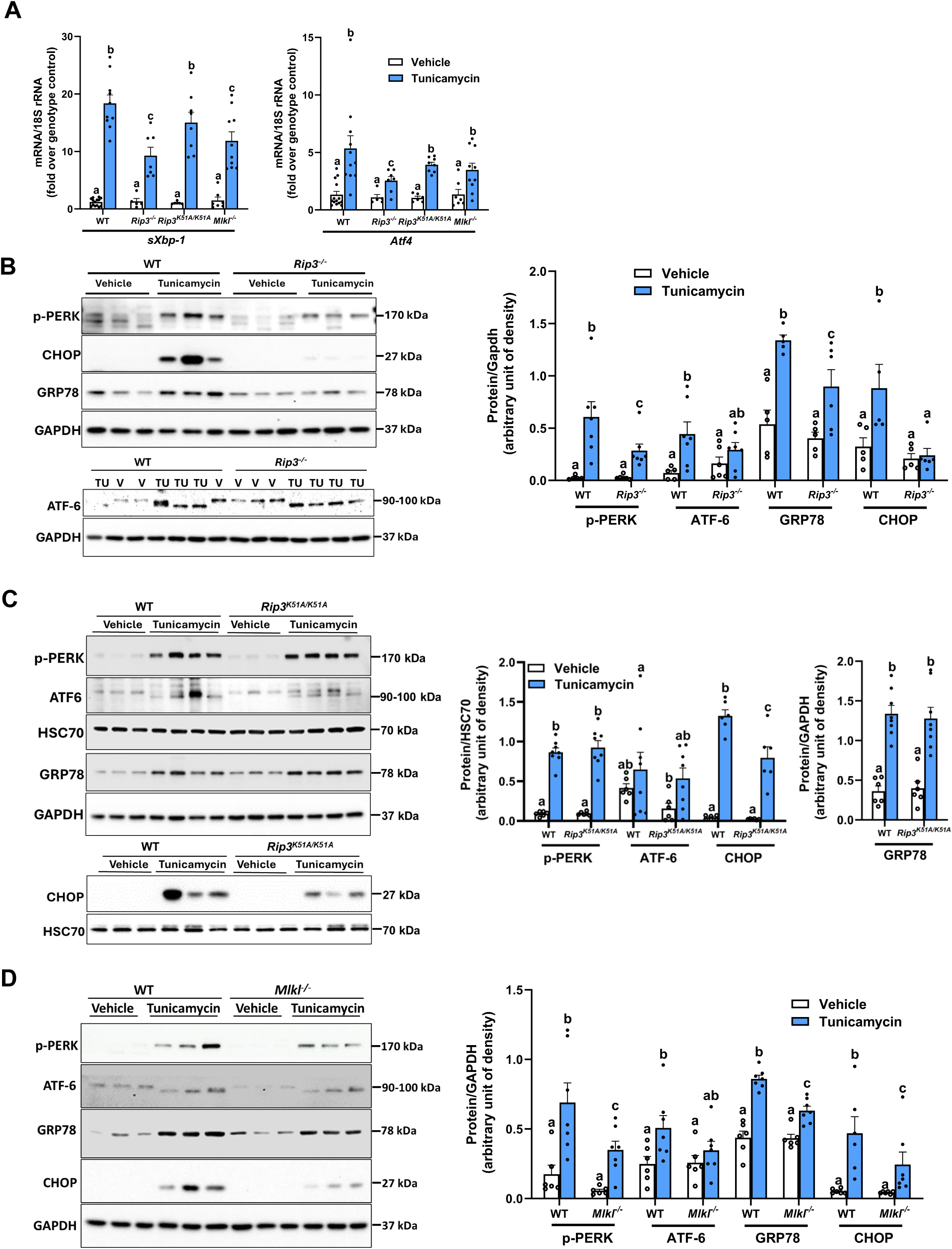
Hepatic ER stress response to tunicamycin is partially impaired in *Rip3^-/-^*, *Rip3^K51A/K51A^*and *Mlkl^-/-^* mice. WT, *Rip3^-/-^*, *Rip3^K51A/K51A^*, and *Mlkl^-/-^* mice were injected with either vehicle or tunicamycin (0.5 mg/kg) for 18 hours. **A.** Expression of *sXbp1* and *Atf4* mRNA was detected in liver of all genotypes using qRT-PCR and normalized to 18S rRNA. **B**. *Rip3^-/-^*, **C.** *Rip3^K51A/K51A^*, and **D.** *Mlkl^-/-^*: Expression of phospho-PERK, GRP78, ATF6, and CHOP was assessed in liver by western blot, densities were quantified using ImageJ and normalized to GAPDH or HSC70; n= 5–8. Blots are processed in parallel for samples from the same experiment. Cropped blots are shown for clarity. Original blots are provided in Supplementary file “*Original western blot images*”. Values represent mean ± SEM. Values with different alphabetical superscripts are significantly different from each other. p < 0.05, assessed by ANOVA.

**Figure 4.**
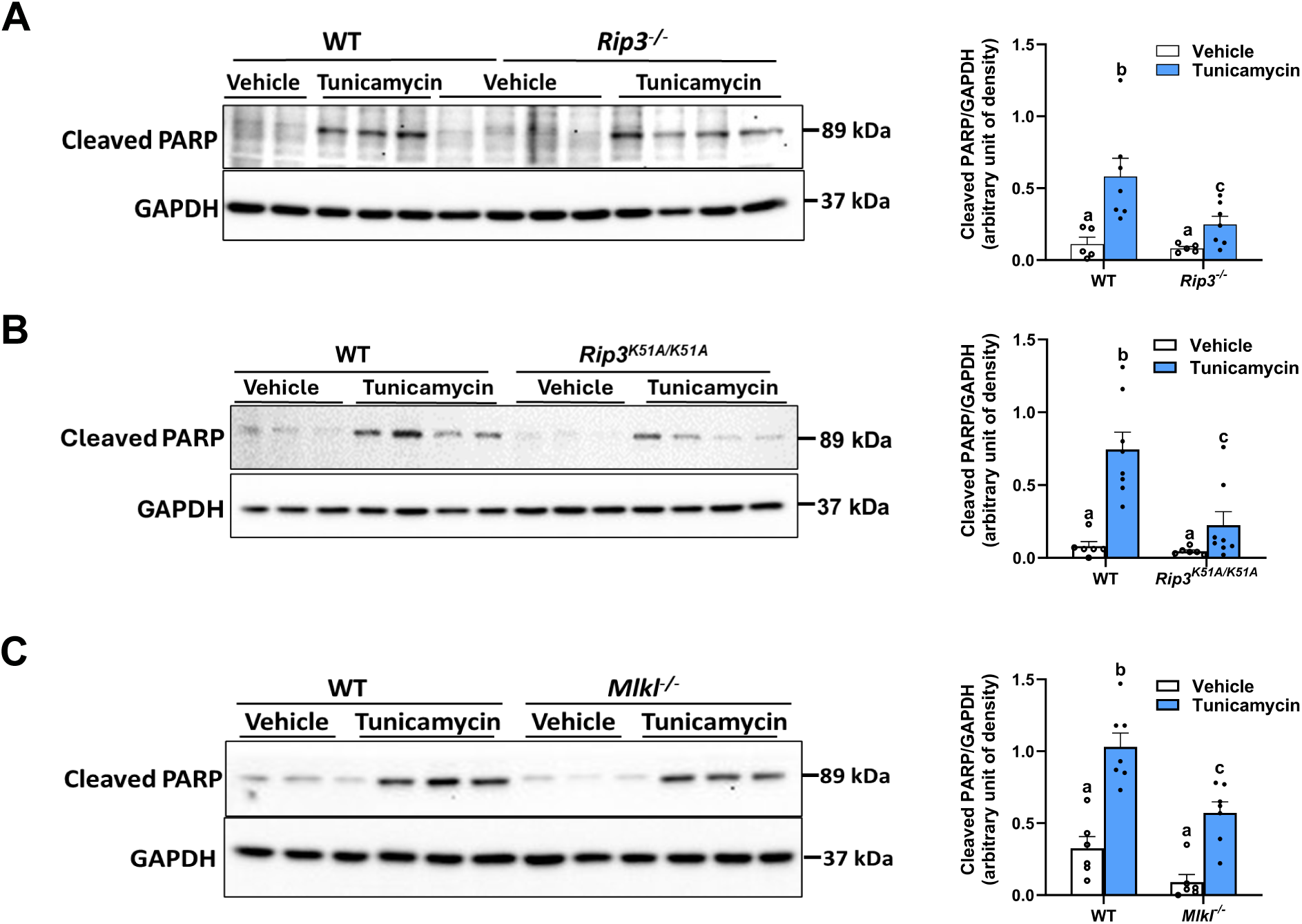
RIP3, MLKL, and RIP3 kinase activity regulate PARP cleavage in the liver. WT, *Rip3^-/-^*, *Rip3^K51A/K51A^*, and *Mlkl^-/-^* mice were injected with either vehicle or tunicamycin (0.5 mg/kg) for 18 hours. **A**. *Rip3^-/-^*, **B.** *Rip3^K51A/K51A^*, and **C.** *Mlkl^-/-^*: Expression of Cleaved PARP was assessed in liver by western blot, densities were quantified using ImageJ and normalized to GAPDH; n= 3-4. Blots are processed in parallel for samples from the same experiment. Cropped blots are shown for clarity. Original blots are provided in Supplementary file “*Original western blot images*”. Values represent mean ± SEM. Values with different alphabetical superscripts are significantly different from each other. p < 0.05, assessed by ANOVA.

To determine whether RIP3 kinase activity contributes to the regulation of tunicamycin-induced ER stress, we examined the RIP3 kinase deficient *Rip3^K51A/K51A^* mice. *Rip3^K51A/K51A^* mice exhibited comparable mRNA expression of ER stress markers to WT controls **(Fig. 3A)**, but there was a marked reduction in accumulation of CHOP protein. Tunicamycin-induced expression of other ER stress proteins was similar between WT and *Rip3^K51A/K51A^*mice (**Fig. 3C**). These results suggest that RIP3 kinase activity selectively regulates CHOP protein expression without altering transcriptional activation of ER stress markers.

Since MLKL acts downstream of RIP3, we further assessed its role in tunicamycin-induced ER stress. Our findings revealed that the mRNA expression of ER stress markers was similarly elevated in *Mlkl^-/-^*and WT mice following treatment **(Fig. 3A)**. However, at the protein level, upregulation of phospho-PERK, GRP78, and CHOP was reduced in *Mlkl^-/-^*compared to WT mice, while ATF6 levels were similar between genotypes (**Fig. 3D**). These findings suggest that MLKL contributes to post-transcriptional regulation of ER stress markers.

### RIP3, MLKL, and RIP3 kinase activity modulate PARP cleavage in the liver

CHOP functions as a key pro-apoptotic factor in the ER stress response, and its suppression may decrease cell death. Since *Rip3^-/-^*, *Rip3^K51A/K51A^* and *Mlkl^-/-^* mice showed reduced ER stress, with particularly lower CHOP expression, after tunicamycin treatment, we investigated whether this reduction protects liver from ER stress-induced apoptosis. Western blot analysis showed that tunicamycin treatment increased the cleavage of PARP, a biochemical hallmark of apoptosis, in WT mice indicating an elevated apoptotic response. In contrast, PARP cleavage was reduced in *Rip3^-/-^*, *Rip3^K51A/K51A^* and *Mlkl^-/-^* mice compared to WT, suggesting reduced apoptosis in response to ER stress **(Fig. 4A-C)**. These findings demonstrate that *Rip3* or *Mlkl* deficiency, or loss of RIP3 kinase activity, diminished ER stress-driven apoptosis in the liver. This reveals a functional link between ER stress pathways and cell death, with RIP3 and MLKL acting as mediators of hepatic apoptosis under conditions of severe ER stress.

### Pharmacological inhibition of RIP3 and MLKL prevents thapsigargin induced ER stress

To delve deeper into the underlying role of RIP3 and MLKL in ER stress regulation, primary hepatocyte cultures were treated with thapsigargin, a potent ER stress inducer. Thapsigargin inhibits the sarco/endoplasmic reticulum Ca² ATPase (SERCA), disrupting calcium homeostasis, leading to the accumulation of misfolded proteins in the ER and subsequent ER stress. While our previous *in vivo* experiments used tunicamycin, we chose thapsigargin for our cell culture studies to explore the contribution of RIP3 and MLKL under a mechanistically distinct model of ER stress induction.

Primary hepatocytes isolated from WT mice were pretreated with either GSK872 (RIP3 kinase inhibitor) or GW806742X (MLKL inhibitor) for 2 hours and then challenged with thapsigargin for 6 hours. Thapsigargin treatment led to a marked increase in the mRNA expression of *sXbp1*, *Atf4*, *Grp78*, and *Chop* compared to vehicle-treated controls. Notably, inhibition of RIP3 or MLKL reduced the expression of *Atf4*, *Grp78*, and *Chop*, while *sXbp1* levels remained unaffected (**Fig. 5A**). At the protein level, thapsigargin increased the expression of phospho-IRE1α and CHOP. This induction was effectively suppressed by pharmacological inhibition of either RIP3 or MLKL (**Fig. 5B and C**). These findings suggest a potential role for RIP3 kinase activity and MLKL in modulating ER stress responses in hepatocytes.

**Figure 5.**
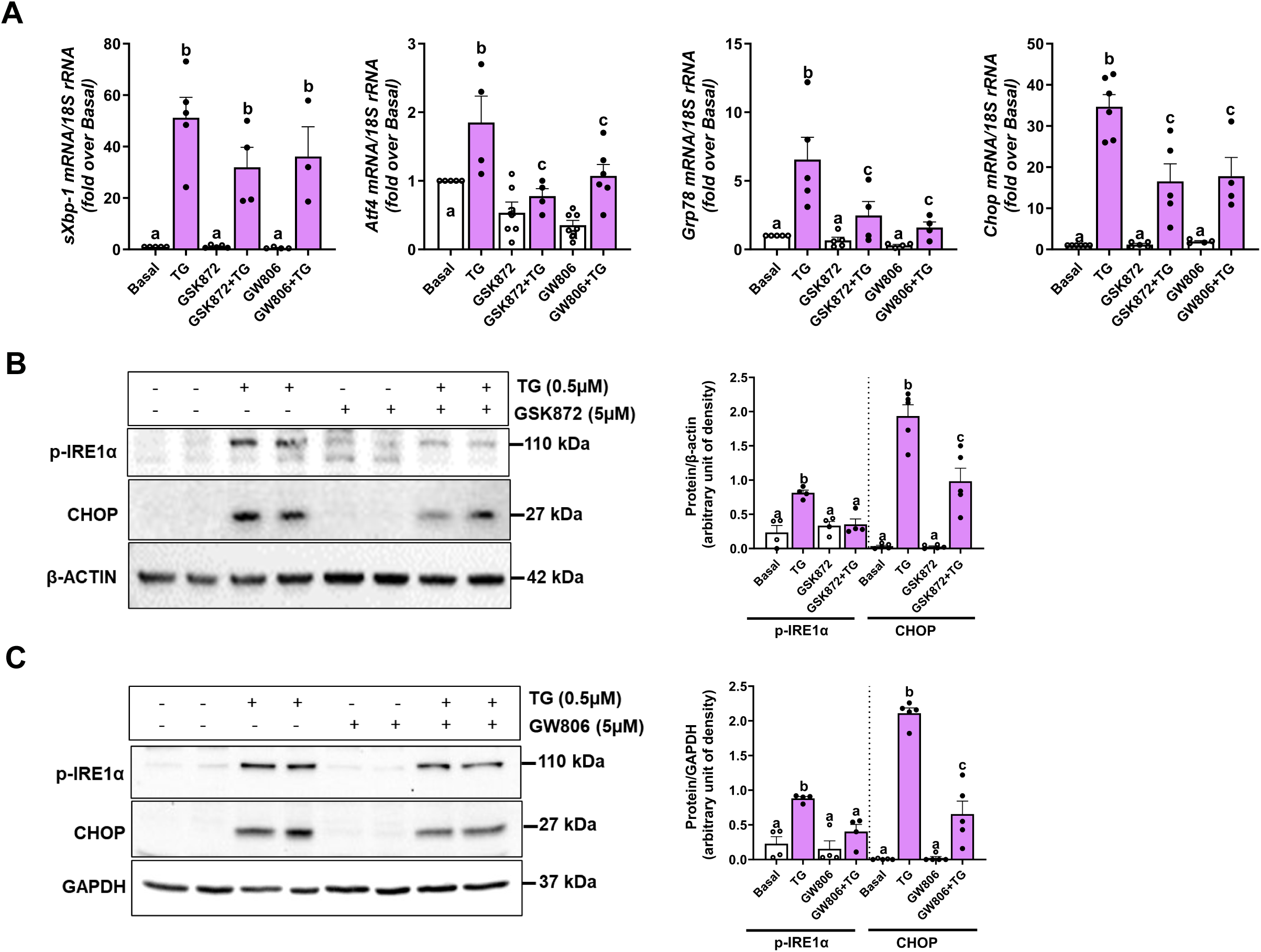
Pharmacological inhibition of RIP3 and MLKL prevents thapsigargin-induced ER stress. Primary hepatocytes from WT mice were pretreated with either GSK872 (RIP3 kinase inhibitor) or GW806742X (MLKL inhibitor) for 2 hours and then treated with 0.5µM thapsigargin (TG) for 6 hours. **A.** Expression of *sXbp1, Atf4, Grp78* and *Chop* mRNA was measured by qRT-PCR and normalized to 18S rRNA. **B**. GSK872 treatment and **C.** GW806742X treatment: Expression of phospho-IRE1α and CHOP was assessed in whole cell lysates by western blot, densities were quantified using ImageJ and normalized to β-Actin or GAPDH; n= 3–7 per group. Blots are processed in parallel for samples from the same experiment. Cropped blots are shown for clarity. Original blots are provided in Supplementary file “*Original western blot images*”. Values represent mean ± SEM. Values with different alphabetical superscripts are significantly different from each other. p < 0.05, assessed by ANOVA.

### Genetic analysis reveals complex interplay between necroptotic mediators and ER stress signaling

Pharmacological inhibitors are valuable tools in elucidating cellular pathways, but their use can be complicated by off-target effects. To circumvent potential confounding effects of inhibitors and to precisely delineate the role of RIP3, its kinase domain and MLKL in regulating thapsigargin-induced ER stress, we employed a genetic approach using primary hepatocytes isolated from WT, *Rip3^-/-^*, *Rip3^K51A/K51A^*and *Mlkl*^-/-^ mice. Following 6-hours thapsigargin treatment, expression of mRNA for ER stress markers (*Atf4*, *sXbp1*, *Grp78*, and *Chop*) were elevated in WT hepatocytes. Compared to WT, *Rip3^K51A/K51A^*hepatocytes showed decreased mRNA expression of *Atf4, Grp78, and Chop* (**Fig. 6A)**. *Mlkl*^-/-^ hepatocytes showed reduced *Grp78* and *Chop* expression, while *Atf4* levels were similar to WT (**Fig. 6C)**. In *Rip3^-/-^*hepatocytes, *Chop* expression was selectively decreased (**Fig. 6A)**. Notably, *sXbp1* levels remained constant across all genotypes (**Fig. 6A and C**). Western blot analysis demonstrated decreased CHOP and phosphorylated IRE1α protein levels in *Rip3^-/-^*, *Rip3^K51A/K51A^* and *Mlkl*^-/-^ hepatocytes compared to WT (**Fig. 6B and D**). These findings are consistent with the impact of pharmacological inhibitors of RIP3 and MLKL and suggest that RIP3 and MLKL have differential effects on the PERK and IRE1α pathways.

**Figure 6.**
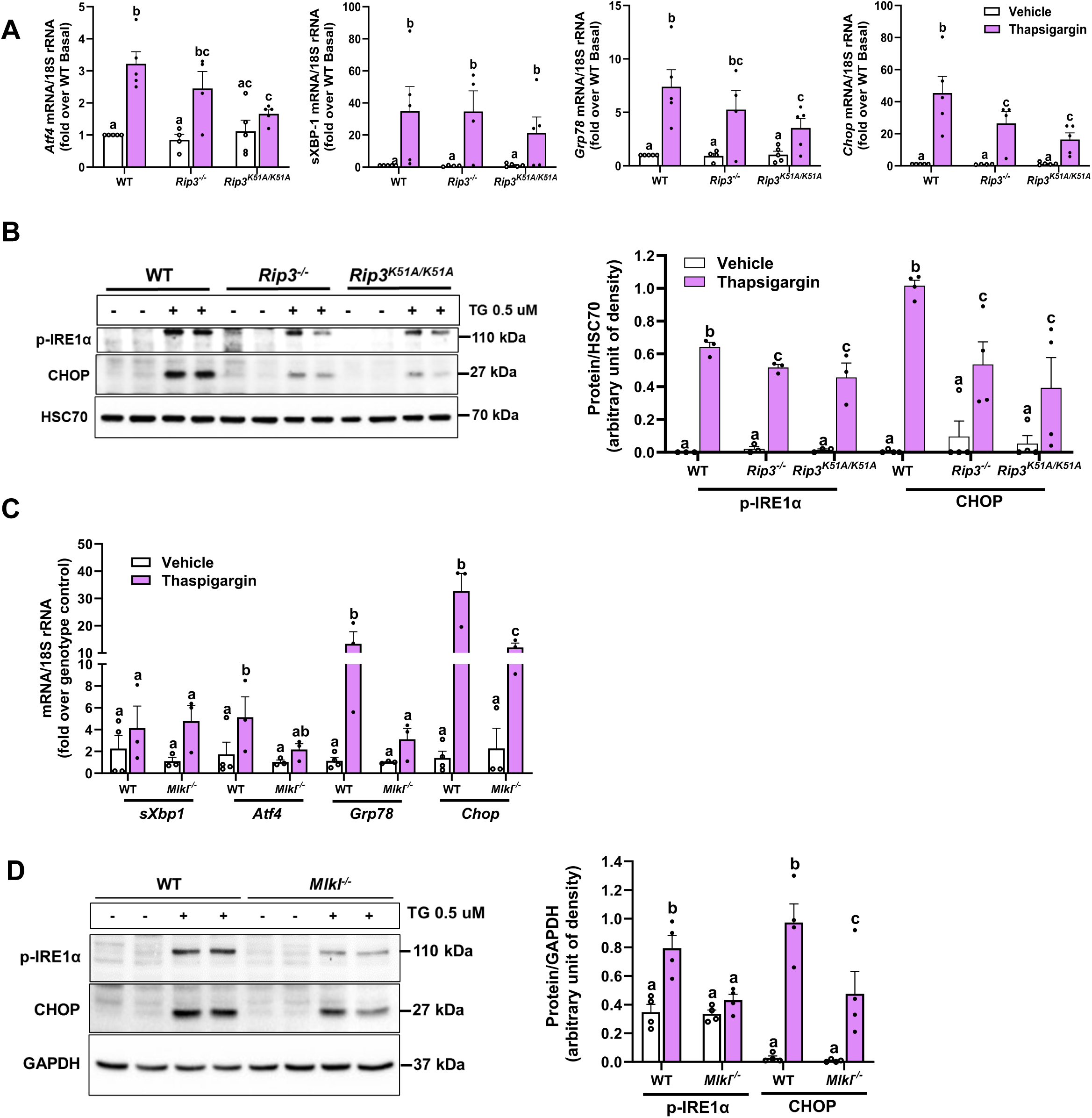
Primary hepatocytes from *Rip3^-/-^*, *Rip3^K51A/K51A^* and *Mlkl^-/-^* mice show attenuated ER stress response to thapsigargin. Primary hepatocytes isolated from WT, *Rip3^K51A/K51A^*, *Rip3^-/-^*, and *Mlkl*^-/-^ mice were exposed to 0.5µM thapsigargin for 6 hours. **A**. *Rip3^-/-^* and *Rip3^K51A/K51A^* **C**. *Mlkl^-/-^*: Expression of *sXbp1, Atf4, Grp78,* and *Chop* mRNA were measured by qRT-PCR and normalized to 18S rRNA. **B**. *Rip3^K51A/K51A^* and *Rip3^-/-^*, and **D.** *Mlkl^-/-^*: Expression of phospho-IRE1α and CHOP was measured in cell lysates by western blot; densities were quantified using ImageJ and normalized to GAPDH or HSC70; n= 3-6 per group. Blots are processed in parallel for samples from the same experiment. Cropped blots are shown for clarity. Original blots are provided in Supplementary file “*Original western blot images*”. Values represent mean ± SEM. Values with different alphabetical superscripts are significantly different from each other. p < 0.05, assessed by ANOVA.

### Deletion of *Rip3* or *Mlkl* protects against ER stress-driven cytotoxicity and facilitates ER expansion in hepatocytes

We investigated the roles of RIP3, its kinase activity, and MLKL in ER stress regulation and hepatocyte survival by isolating primary hepatocytes from WT, *Rip3^-/-^*, *Rip3^K51A/K51A^* and *Mlkl*^-/-^ mice and treating them with thapsigargin for 24 hours. Cytotoxicity analysis showed that *Rip3^-/-^*, *Rip3^K51A/K51A^*and *Mlkl*^-/-^ hepatocytes were protected from thapsigargin-induced cell death, compared to WT, suggesting that RIP3 and MLKL contribute to ER stress-mediated cytotoxicity (**Fig. 7A**).

**Figure 7.**
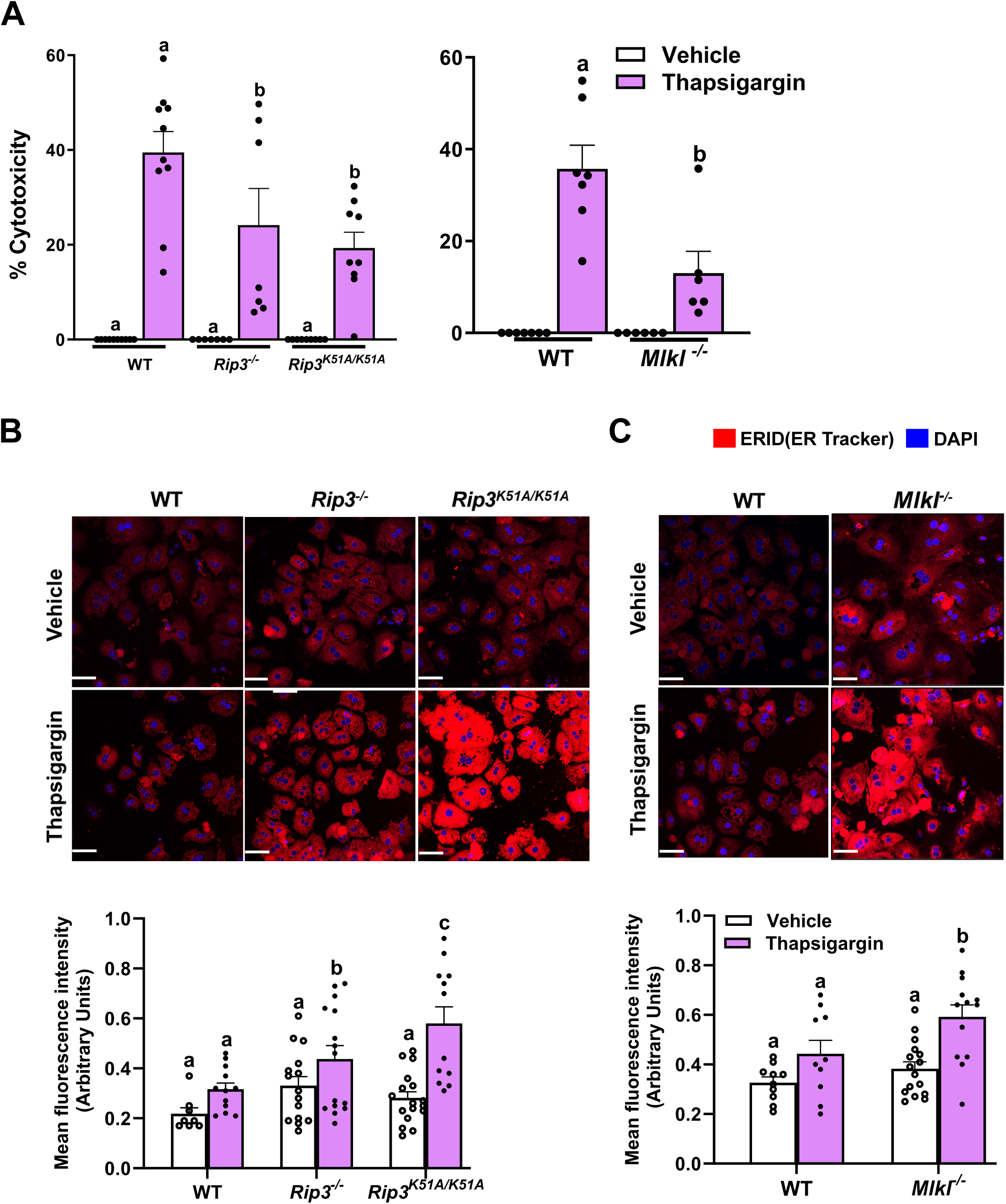
Genetic ablation of *Rip3*, RIP3 kinase activity or *Mlkl* protects hepatocytes from thapsigargin-induced toxicity and promotes ER expansion. Primary hepatocytes isolated from WT, *Rip3^-/-^*, *Rip3^K51A/K51A^*, and *Mlkl*^-/-^ mice were treated with 0.5µM thapsigargin for 24 hours for MTS assay and 16 hours for ER staining. **A.** MTS assay was performed to determine thapsigargin-induced hepatotoxicity, n = 6–12 per group. **B-C.** For ER morphology primary hepatocytes from **B**. WT, *Rip3^-/-^*, and *Rip3^K51A/K51A^* and **C**. WT and *Mlkl*^-/-^ grown on collagen-precoated Permanox™ plastic chamber slides were fixed and stained with the ER marker (ER ID dye; red) and mounted with mounting media with DAPI (blue). Cells were captured using a 40X oil lens, individual data points on graph represent individual values from triplicate images from 3-4 different experiments. Scale bars: 50 μm. Values represent mean ± SEM. Values with different alphabetical superscripts are significantly different from each other. p < 0.05, assessed by t test or ANOVA.

To elucidate the mechanisms underlying hepatocyte resistance to ER stress-induced cytotoxicity, we assessed ER membrane expansion using ER-ID (an endoplasmic reticulum-selective dye) staining. ER expansion is recognized as a protective adaptation that increases the organelle’s capacity to handle protein-folding stress, thereby mitigating ER stress and potentially reducing susceptibility to cell death. Following thapsigargin challenge, hepatocytes from *Rip3^-/-^*, *Rip3^K51A/K51A^* and *Mlkl*^-/-^ mice displayed pronounced ER expansion, in contrast to the lower expansion observed in WT cells (**Fig. 7B–C**). These findings suggest that the deficiency of *Rip3,* RIP3 kinase activity or *Mlkl* enhances ER biogenesis, supporting cellular survival under ER stress conditions. Collectively, this highlights a key role for RIP3 and MLKL in limiting the adaptive response to ER stress and promoting cell death.

### Deficiency of *Rip3*, RIP3 kinase activity and *Mlkl* enhances ER sheet formation and adaptive remodeling during ER stress

To further investigate structural adaptations of the ER that contribute to hepatocyte resistance against ER stress-induced cytotoxicity, we analyzed ER morphology using ER-ID staining and three-dimensional reconstruction of Z-stacks images in primary hepatocytes isolated from WT, *Rip3^-/-^*, *Rip3^K51A/K51A^* and *Mlkl*^-/-^ mice under both vehicle and thapsigargin-treated conditions (**Fig. 8A-D**). Striking differences in ER architecture were observed across genotypes: while WT hepatocytes exhibited a predominantly tubular ER network, characterized by thin, interconnected structures distributed across the periphery of the cell, hepatocytes from *Rip3^-/-^*, *Rip3^K51A/K51A^* and *Mlkl*^-/-^ mice displayed a higher proportion of sheet-like ER structures even under basal conditions, indicating intrinsic alterations in ER organization in the absence of these necroptosis modulators.

**Figure 8.**
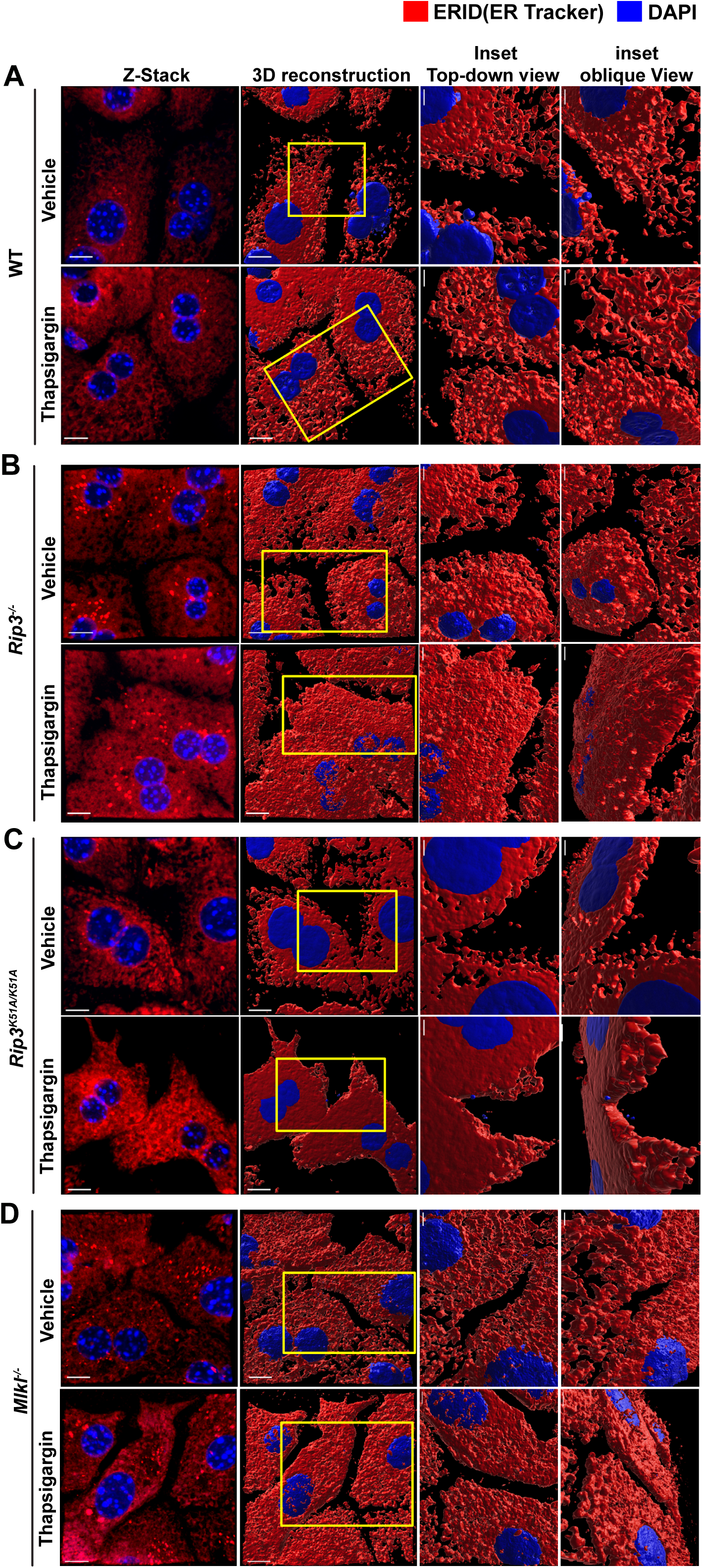
Enhanced ER sheet formation and adaptive morphological changes in primary hepatocytes from *Rip3^-/-^*, *Rip3^K51A/K51A^*, and *Mlkl^-/-^* mice during ER stress. Primary hepatocytes from WT, *Rip3^-/-^*, *Rip3^K51A/K51A^* and *Mlkl*^-/-^ mice were treated with vehicle or thapsigargin (0.5 μM, 16 hours) to induce ER stress as described in Figure 7. Cells were stained with ER-ID and captured in z-stacks using a 40× oil immersion objective with 4× zoom (n=3-4 per condition). 3D reconstructions of z-stacks were generated using IMARIS software, and the *Surfaces* module was employed to visualize nuclei and ER structures. (**A-D**) Representative z-stack projections and 3D reconstructions showing ER morphology in **A**. WT, B. *Rip3^-/-^*, **C**. *Rip3^K51A/K51A^*, and **D**. *Mlkl^-/-^* primary hepatocytes under basal and stress conditions. Insets display top-down and oblique views of selected regions at higher magnification to highlight detailed ER architectural features. Scale bars: 10 μm (main images); 4 μm (insets).

Upon thapsigargin-induced ER stress, WT hepatocytes maintained a predominantly tubular ER architecture with only modest expansion. In contrast, hepatocytes from *Rip3^-/-^*, *Rip3^K51A/K51A^* and *Mlkl*^-/-^ mice exhibited extensive reorganization of the ER network, with more prominent sheet formation. These differential sheet–tubule rearrangements are especially evident in the 3D reconstructions when examined from different angles (**Fig. 8A–D**, top-down and oblique view insets). The structural differences in ER morphology across different genotypes and treatment conditions are further highlighted in the animation (rotational videos) of 3D reconstructed images from Figure 8 (**Supplemental videos 1–8**).

Collectively, these findings suggest that RIP3 and MLKL act as negative regulators of adaptive ER remodeling during stress, and that increased ratio ER sheets to tubules may support greater protein folding capacity, thereby promoting cellular homeostasis and protecting hepatocytes from ER stress-induced cell death.

## Discussion

Our study provides novel insights into the complex interplay between necroptotic mediators and ER stress signaling in the context of ALD and pharmacologically-induced ER stress. The findings demonstrate that RIP3, RIP3 kinase activity, and MLKL play crucial roles in modulating hepatic ER stress responses and cell death pathways, with potential implications for the pathogenesis and treatment of ALD and other ER stress-related liver diseases.

The GSEA of RNA-seq data from sAH patients revealed enrichment of ER stress response pathways, aligning with previous studies highlighting the role of ER stress in ALD pathogenesis ^[3, 7, 29, 30]^. This finding suggests that targeting ER stress pathways could be a promising therapeutic approach for ALD. The observation that *Rip3* deficiency attenuates ethanol-induced upregulation of ER stress markers provides evidence for RIP3 involvement in alcohol-mediated ER stress. These observations point to a multifaceted role for RIP3 in regulation of cellular stress, not limited to necroptosis. The time-dependent upregulation of RIP3 and MLKL following tunicamycin treatment suggests an association between ER stress and the induction of necroptotic mediators. This bidirectional relationship suggests a potential feed-forward loop where ER stress induces RIP3 and MLKL expression, which in turn may exacerbate ER stress responses.

A key question is whether the attenuation of ER stress markers in *Rip3^-/-^* and *Mlkl^-/-^*mice reflects a direct role of RIP3/MLKL in the regulation of ER stress or a secondary effect of reduced necroptosis. Three lines of evidence suggest that RIP3/MLKL have a direct role in regulation of ER structure and function: 1) the time course of increased plasma ALT and ER stress activation are similar, suggesting injury and ER stress occur in parallel; 2) *Rip3* or *Mlkl* deficiency selectively blunts PERK and IRE1α but not ATF6 signaling; and 3) hepatocytes lacking *Rip3* or *Mlkl* display intrinsic architectural shifts in the ER, characterized by an increased sheet-to-tubule ratio even under basal conditions when no injury is present. Together, these findings are consistent with the hypothesis that RIP3 and MLKL act as direct modulators of ER stress signaling and morphology, extending their function beyond necroptosis.

Our genetic and pharmacological inhibition studies uncover a non-canonical role for RIP3 and MLKL in modulating hepatic ER stress, independent of their established function in necroptosis. Deficiency of *Rip3*, RIP3 kinase activity, or *Mlkl* partially reduced tunicamycin-induced ER stress, particularly by attenuating the PERK and IRE1α (sXbp1) pathways. While *Rip3^-/-^* mice showed reduced *Atf4* and *sXbp1* mRNA and lower levels of phospho-PERK, GRP78, and CHOP proteins, *Rip3^K51A/K51A^* mice exhibited selective reduction in CHOP protein, indicating a post-transcriptional regulatory role for RIP3 kinase activity. Similarly, *Mlkl* deficiency blunted ER stress protein expression without affecting mRNA levels, consistent with a post-transcriptional regulation of at least some ER stress proteins.

These reductions in ER stress proteins were associated by decreased PARP cleavage, indicating that RIP3 and MLKL promote ER stress-induced apoptosis. Pharmacological and genetic inhibition of RIP3 and MLKL in thapsigargin-treated hepatocytes further supported their roles in amplifying ER stress signaling. Notably, *sXbp1* mRNA levels remained unchanged across genotypes, despite reduced phospho-IRE1α, highlighting a disconnect between transcriptional and post-transcriptional regulation. This discrepancy underscores the importance of evaluating both levels when dissecting ER stress pathways ^[31]^.

A key observation in our study is the impact of RIP3 and MLKL on ER morphology and architecture. Confocal microscopy analysis demonstrated that hepatocytes from *Rip3^-/-^*, *Rip3^K51A/K51A^*, and *Mlkl^-/-^* mice exhibited enhanced ER expansion in response to thapsigargin, in contrast to WT hepatocytes, which showed minimal ER expansion. This suggests that RIP3 and MLKL may actively suppress ER adaptation, potentially exacerbating ER stress and promoting hepatocyte death. The ability of ER expansion to mitigate ER stress has been reported in previous studies, wherein increased volume of the ER correlates with improved protein folding and stress resolution ^[32–37]^. Given that unresolved ER stress leads to cell death via apoptosis and/or necroptosis, the absence of RIP3 or MLKL may allow hepatocytes to better adapt to stress conditions, thereby enhancing their survival.

While ER enlargement was evident, the more remarkable insight was how RIP3 and MLKL orchestrated the architectural organization and structural dynamics of the ER. Using ER-ID staining and three-dimensional confocal reconstruction, we found that hepatocytes deficient in *Rip3*, RIP3 kinase activity, or *Mlkl* exhibited an intrinsic shift from a tubular ER network to a greater proportion of sheet-like structures, even under basal conditions. Under thapsigargin-induced ER stress, this phenotype became even more pronounced: while WT hepatocytes maintained a tubular ER architecture with only modest expansion, *Rip3^-/-^*, *Rip3^K51A/K51A^*, and *Mlkl^-/-^* hepatocytes showed extensive ER reorganization with prominent sheet formation, as clearly visualized in 3D reconstructions insets (**Fig. 8A-D**).

These morphological adaptations likely confer a survival advantage, as ER sheet structures are known to enhance protein folding capacity and promote adaptive stress responses. In line with previous studies ^[32, 38]^, our findings suggest that increased ER sheet formation may serve as a compensatory mechanism to maintain proteostasis during ER stress. Thus, RIP3 and MLKL appear to act as negative regulators of adaptive ER remodeling, limiting the capacity of hepatocytes to withstand ER stress.

While our analyses provide valuable insights into ER structural remodeling, it is important to recognize the methodological limitations. Confocal imaging lacks the resolution to capture finer ultrastructural differences, such as distinguishing between subtle variations in sheet and tubular networks or examining ER-mitochondrial contact sites. Future studies could benefit from employing correlative electron microscopy or advanced super-resolution imaging, which would enable more precise visualization of ER ultrastructure and validation of the observed changes ^[34]^. Furthermore, integrating lipidomic and proteomic analyses may clarify how RIP3 kinase activity regulates ER membrane composition to influence structural plasticity during stress. Addressing these questions will advance our understanding of ER architecture dynamics in ALD and may reveal new therapeutic targets to enhance ER stress resilience.

The observation that loss of *Rip3*, its kinase activity or *Mlkl* attenuates ER stress-induced apoptosis, as evidenced by reduced cleaved PARP levels in liver, further supports their role in promoting cell death under conditions of severe ER stress. Our results suggest that targeting the RIP3-MLKL axis could be a promising strategy for mitigating ER stress-induced liver injury in ALD and other liver diseases characterized by chronic ER stress.

In conclusion, our study unveils a novel role for RIP3 and MLKL in regulating hepatic ER stress responses, highlighting their impact on ER stress pathways and the pathogenesis of ALD. This interplay between RIP3-MLKL axis and ER stress response underscores the need for further research to elucidate the molecular mechanisms governing these interactions and their implications for liver health and disease. Future studies employing high-resolution imaging techniques and integrative -omics approaches will be essential to uncover the precise structural and molecular changes driving enhanced stress resilience in *Rip3/Mlkl*-deficient hepatocytes. By deepening our understanding of ER architecture dynamics and stress adaptation mechanisms, this work lays the foundation for developing targeted interventions to modulate ER stress resilience and improve liver health in ALD and related disorders.

## Abbreviations

ALD: alcohol-associated liver disease;
ALT: alanine aminotransferase;
AST: aspartate aminotransferase;
ATF4: activating transcription factor 4;
ATF6: activating transcription factor 6;
CHOP: C/EBP homologous protein;
DAPI: 4′,6-diamidino-2-phenylindole;
GRP78: glucose-regulated protein 78;
GSEA: gene set enrichment analysis;
H&E: hematoxylin and eosin;
HSC70: heat shock cognate 70;
IRE1α: inositol-requiring enzyme 1 alpha;
MTS: 3-(4,5-dimethylthiazol-2-yl)-5-(3-carboxymethoxyphenyl)-2-(4-sulfophenyl)-2H-tetrazolium (cell viability assay);
NES: normalized enrichment score;
PARP: poly (ADP-ribose) polymerase;
PBS: phosphate-buffered saline;
PERK: protein kinase RNA-like ER kinase;
PVDF: polyvinylidene fluoride;
qRT-PCR: quantitative real-time reverse transcription polymerase chain reaction;
sXBP1: spliced X-box binding protein 1;
TG: thapsigargin;
TU: Tunicamycin;
UPR: unfolded protein response.

## Acknowledgments

The authors sincerely thank Dr Zhaoli Sun at Johns Hopkins University for the human liver specimens. The authors thank Amit Gupta, PhD (Genomic Medicine Institute, Cleveland Clinic) for his assistance in revising Figure 1. The authors also acknowledge the assistance of the Cleveland Clinic Lerner Research Institute Imaging Core in providing microscopy services. This work was also supported by the Harrington Physician-Scientist Pathway of University Hospitals / Case Western Reserve University (to JT). Illustrations (Graphical abstract) used in this publication was created by David Schumick, BS, CMI, and are reprinted with the permission of the Cleveland Clinic Enterprise Creative Services ©2025. All Rights Reserved.

## Declaration of generative AI and AI-assisted technologies in the writing process

*During the preparation of a few paragraphs of Introduction and Discussion part of this work the first author (RKA) used ChatGPT in order to improve language and readability only. After using this ChatGPT, the author(s) reviewed and edited the content as needed and takes full responsibility for the content of the publication*.

