## Supplementary figures and images for "RIP3 and MLKL regulate Hepatic ER stress in alcohol-associated liver disease and pharmacological ER stress models: insights beyond necroptosis"

### Supplemental fig 1

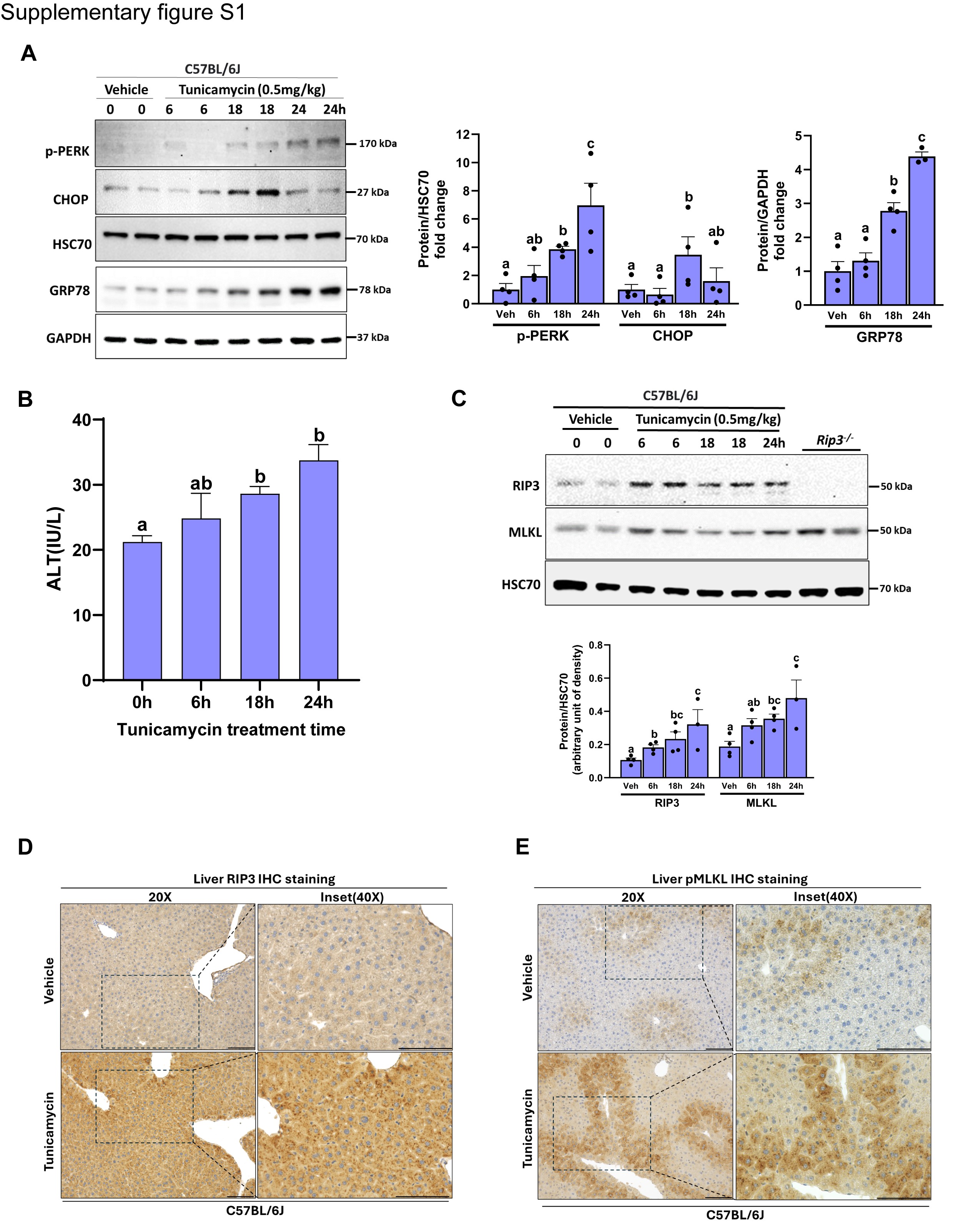

### Supplemental Fig 2

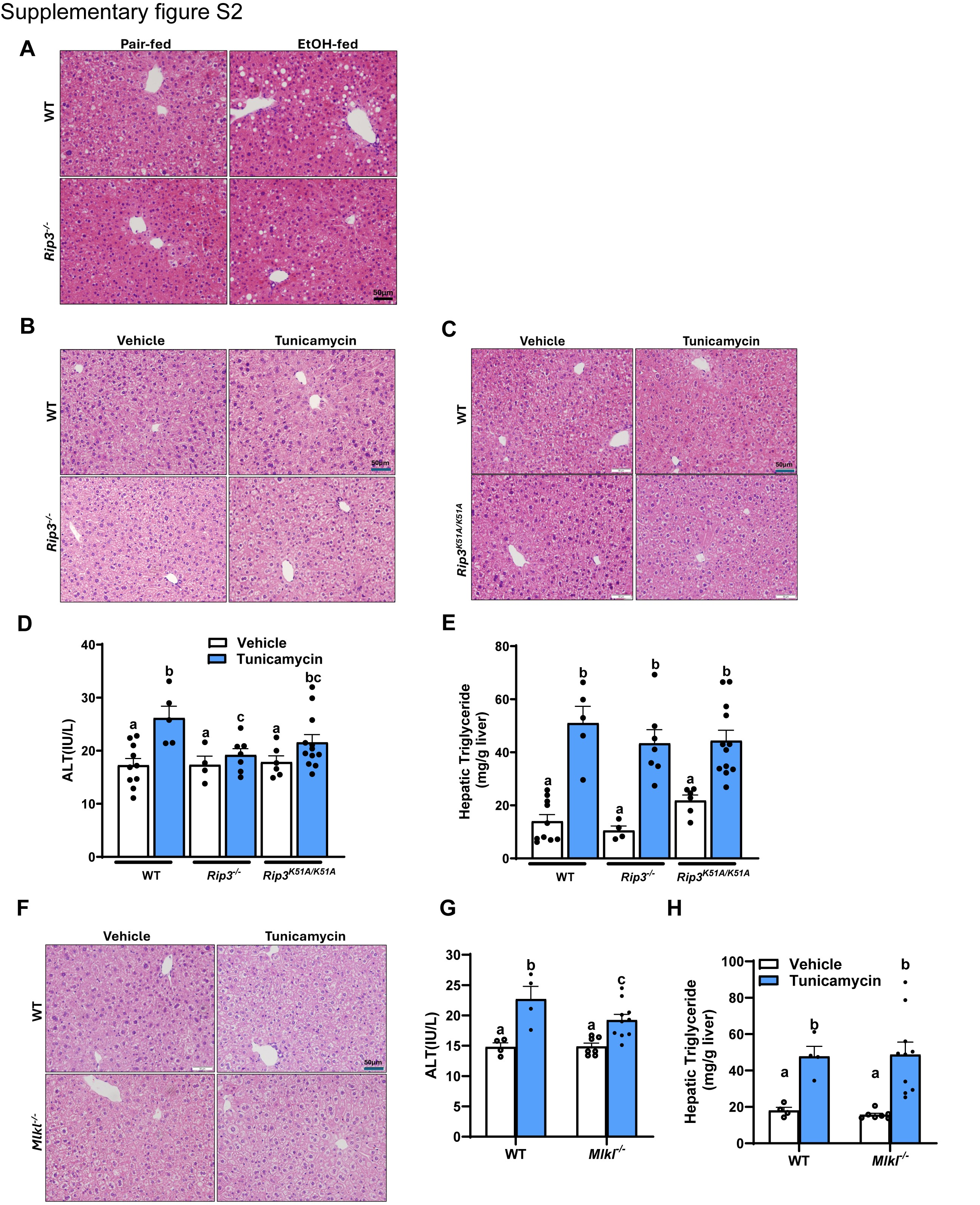
